# The Limits of Haplotype-Based Approaches: Exploring the Applicability of the Li and Stephens Haplotype-Copying Model to Ancient Samples

**DOI:** 10.1101/2023.06.21.545876

**Authors:** Isabel Díaz-Pinés Cort, Joshua Daniel Rubin, Peter Wad Sackett, Gabriel Renaud

**Affiliations:** Department of Health Technology, Technical University of Denmark, Kongens Lyngby, Denmark; Costerton Biofilm Center, Department of Immunology and Microbiology, University of Copenhagen, Copenhagen, Denmark

## Abstract

The Li and Stephens (LS) haplotype-copying model is a seminal framework that represents a target haplotype as an imperfect mosaic of a set of reference haplotypes. Using a hidden Markov model, it can switch from different source haplotypes to model recombinations. This model has been used in several applications in modern populations including phasing and inference of ancestry. However, recent publications have looked at the applicability of the model to using ancient individuals as targets and modern reference panels as source data. Previous research exploring the impact of time separation between the modern references and the ancient target on the model’s behavior relied on coalescent simulation to generate genetic variation data, which could lead to an underestimation of the ancient population’s genetic diversity. Further, these simulations were restricted to a relatively short time period of anatomically modern human history. To overcome these limitations, our study evaluates the robustness of the LS model using forward-simulated data enabling us to sample haplotypes that do not have direct descendants among the modern population. Additionally, we evaluate the model under the simple demographic scenario of a constant-sized continuous population starting 1.5M years ago to isolate the effect of time separation. Results indicate good performance for target haplotypes up to 900,000 years old, suggesting potential applicability to ancient DNA (aDNA) from anatomically modern humans. Although more complex demographic scenarios should be considered for a definitive answer, this research serves as a starting point for evaluating the haplotype-copying framework in aDNA data analysis.

## 1 Introduction

The Li and Stephens (LS) model, a hidden Markov model, has proven to be a potent instrument for the analysis of genetic data, especially in understanding haplotype inference and recombination events (Li and Stephens, 2003). Its core assumption is that each individual’s haplotype in a sample is a mosaic of the haplotypes of the other individuals, forming the foundation of its “copying” model (Li and Stephens, 2003). The LS model has been extensively utilized in population genetics studies, particularly in the context of modern haplotypes (Stephens et al., 2001). An application of this model within modern haplotypes is Chromopainter, which identifies shared ancestry of haplotypes and characterizes population substructure by building a co-ancestry matrix. Utilizing the LS model, `Chromopainter` “paints” a recipient haplotype using chunks of donor haplotypes, capturing the most relevant genealogical information about a haplotype sample and hypothesizing that the most recent genealogical events sufficiently characterize the current substructure of a population (Lawson et al., 2012). The LS model has also been employed to infer modern haplotypes from ancestral ones, as seen in the implementation of the software `tsinfer` (Kelleher et al., 2019). However, the exploration of inferring ancient haplotypes from modern ones using the LS model is still an active area of research, presenting a promising avenue for further advancements in the field of genetic studies.

Indeed, in recent work, researchers have been using a scenario of the Li and Stephens model to address problems with ancient DNA (aDNA), where the target haplotype corresponds to an ancient individual, and the reference panel comprises modern haplotypes from present-day individuals. Prophaser, one such tool, leverages the LS model to handle genotype likelihoods instead of SNP data, resulting in phased haplotypes and imputation of missing variants (Ausmees and Nettelblad, 2022). Although it outperforms other commonly used pipelines, its applicability to older genomes remains unproven, and its performance at low coverage depths could be overestimated due to testing on down-sampled modern genomes. Another tool, `hapCon`, applies the LS model to estimate contamination rates in ancient male X chromosomes, using three error parameters (Huang and Ringbauer, 2022). The tool has demonstrated good performance even with old samples, but the authors note that genetic similarity between the contamination and endogenous sources can introduce a bias towards the contamination source (Huang and Ringbauer, 2022). Both tools highlight the utility and adaptability of the LS model in studying aDNA, but also underscore the need for rigorous testing in diverse scenarios.

In this study, we are focused on testing the applicability of the Li and Stephens model to ancient DNA samples using modern reference haplotypes. Our aim is to investigate how well this model performs across a simulated population spanning 1.5 million years, or approximately 50,000 generations.

This is a significant undertaking as it offers valuable insight into how genetic relationships between ancient and modern populations can be discerned. These simulations represent perhaps the best-case scenario for human data with a large effective population, continuity and perfect resolution of the alleles.

The first stage of our study involved estimating the Maximum Likelihood Estimators (MLEs) for the jump rate and copying error parameters of the Li and Stephens model. We found that the jump rate MLE showed an increase in relation to the sample age until around 25,000-30,000 generations ago, after which the relationship turned exponential. In contrast, the copying error MLE maintained a consistent linear relationship with sample age throughout the period.

In comparing the optimal path divergence of the Li and Stephens model to that of a baseline model, our findings revealed interesting patterns. While the divergence from the optimal path for the Li and Stephens model showed a linear increase until about 30,000 generations, this trend plateaued thereafter. Notably, the mean divergence from the modern reference haplotypes increased at a higher rate than the divergence from the optimal path. We also observed a slight decrease after 40,000 generations, indicating potential limitations in the model’s performance with older samples.

Finally, the variance of divergences from the reference haplotypes plotted against sample age showed an exponential decrease. This suggests that, over time, the target haplotypes become equally divergent from all modern reference haplotypes. These results underscore the utility of the Li and Stephens model in elucidating relationships between ancient and modern populations, while also highlighting areas for further refinement when dealing with particularly ancient samples.

## 2. Material and Methods

In this section, we briefly summarize the methods employed in this study. A more detailed description can be found in the Supplementary Methods section (1).

### 2.1 Simulation and Data Generation

We performed forward-time simulations to generate genomic data for a population starting 1.5 million years ago under a simple demographic model that represents the best-case scenario under which the haplotype-copying framework could be evaluated.. This model consisted of a single continuous population with constant population size, neutral mutation, and random mating. The simulation parameters were informed by empirical estimates from the literature, including generation time, recombination rate, mutation rate, and effective population size (Tremblay and Vézina, 2000; Gutenkunst et al., 2009; Jensen-Seaman et al., 2004; Renaud et al., 2019). The simulated sequence was 1Mb long.

The genomic data at the population level was simulated using the slendr package in R (Petr, 2022), with the forward-time simulator `SLiM` as a backend engine (Haller et al., 2019). The output consisted of tree-sequence files, which were recapitulated using `slendr` and `msprime` (Baumdicker et al., 2022).

We created a reference panel of size *k* = 100 modern haplotypes and uniformly sampled target haplotypes at random every 200 generations. The haplotype-copying model, parameter estimation, Viterbi decoding, and model evaluation were performed following the methodology described in the Supplementary Methods section (1).

### 2.2 Parameter Estimation and Model Evaluation

For the estimation of model parameters and evaluation of the haplotype-copying model, we employed the following procedures:

#### 2.2.1 Maximum Likelihood Estimation

We obtained Maximum Likelihood Estimates (MLEs) of the jump rate and copying error parameters, denoted 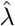 and 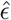 respectively, via direct maximization of the likelihood, as described in Supplementary Section 1.4.4.

The implementation followed the work by Biddanda *et al*. (Biddanda et al., 2022). The MLEs were first computed individually, and then jointly estimated using two-dimensional numerical minimization. The standard errors for the MLEs were obtained using a finite-difference approximation to the second derivative of the joint log-likelihood surface.

#### 2.2.2 Viterbi Decoding

After obtaining the MLEs of the jump rate and copying error parameters, we used the Viterbi algorithm to compute the most probable path of hidden states and its log posterior probability for each test haplotype, as detailed in Supplementary Section 1.4.5. We also applied Viterbi decoding to the data using a baseline model, where all the transition probabilities are equal to 1*/k*, and the copying error parameter is 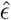

#### 2.2.3 Model Evaluation Metrics

To evaluate the model’s performance as sample age increases, we employed three metrics: the MLE of the jump rate parameter 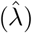), the divergence between the target haplotype and the optimal path of hidden states, and the log posterior probability of the optimal path, as explained in Supplementary Section 1.4.6. We compared the divergence and log posterior probability obtained under the maximum likelihood (ML) model to those obtained under the baseline model.

Additionally, we analyzed the distribution of divergences from the modern reference haplotypes to the target haplotypes as a function of their age to provide insight into the model’s behavior as the target haplotypes’ age increases, as discussed in Supplementary Section 1.4.6.

All code and data used in this project can be found in the GitHub repository at https://github.com/isadpc/HapCopying. For further information on the code and data, refer to Section 1.4.7.

## 3. Results

In this section, we present the main results of our research.

### 3.1 MLEs

MLEs of the jump rate and copying error parameters were obtained. As can be seen in Figure 1a, the MLE of the jump rate parameter, 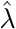, increases monotonically with sample age. The trend is approximately linear until a critical point (around 25,000-30,000 generations ago) after which the relationship with sample age becomes exponential. On the other hand, the copying error MLE, 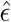 maintains a positive linear relationship with sample age throughout (1b).

**Figure 1:**
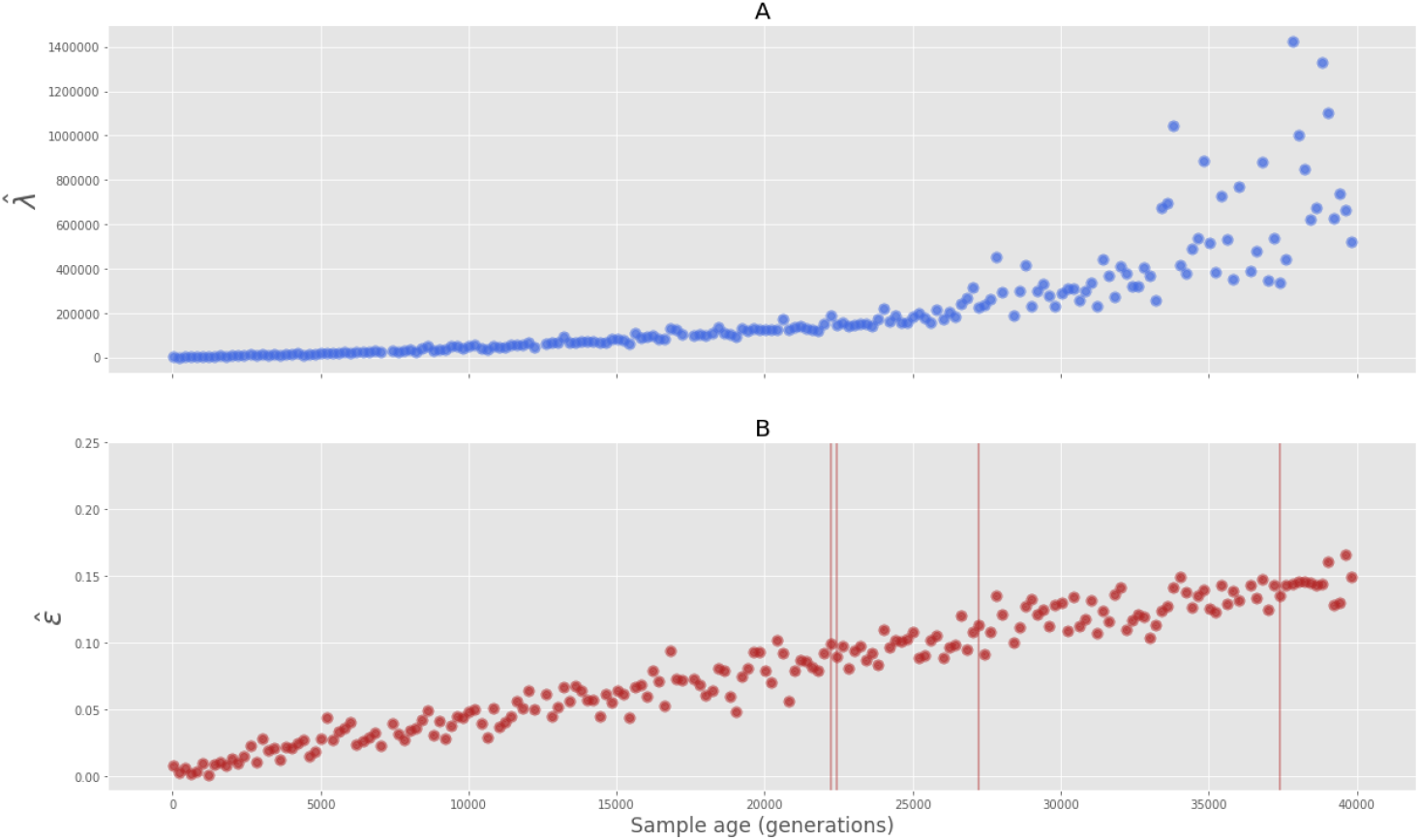
MLEs versus sample age. **A** jump rate MLE. **B** copying error MLE. Only results until 40,000 generations back in time are shown for the trends to be visible. The vertical lines correspond to the error bars.

### 3.2 Comparison to a Baseline Model

After estimating the model parameters, the optimal path of hidden states and its log posterior probability were obtained via the Viterbi algorithm. Again, the divergences from the target haplotypes to the optimal paths were computed at each time point. The same was done for the baseline model, where jumping to any reference haplotype or staying in the current one is equiprobable.

For the ML model, the divergence from the optimal path increases linearly with sample age until about 30,000 generations ago, where a slight decrease follows a plateau (Figure 2a). In the case of the baseline model, the divergence from the optimal path increases monotonically with sample age, again and as expected, at a slower rate than for the ML model (Figure 2b). Note that since all transitions are equiprobable for the latter model, the divergence from the optimal path reflects the number of positions at which the ancient target haplotype differs from all the modern reference haplotypes.

**Figure 2:**
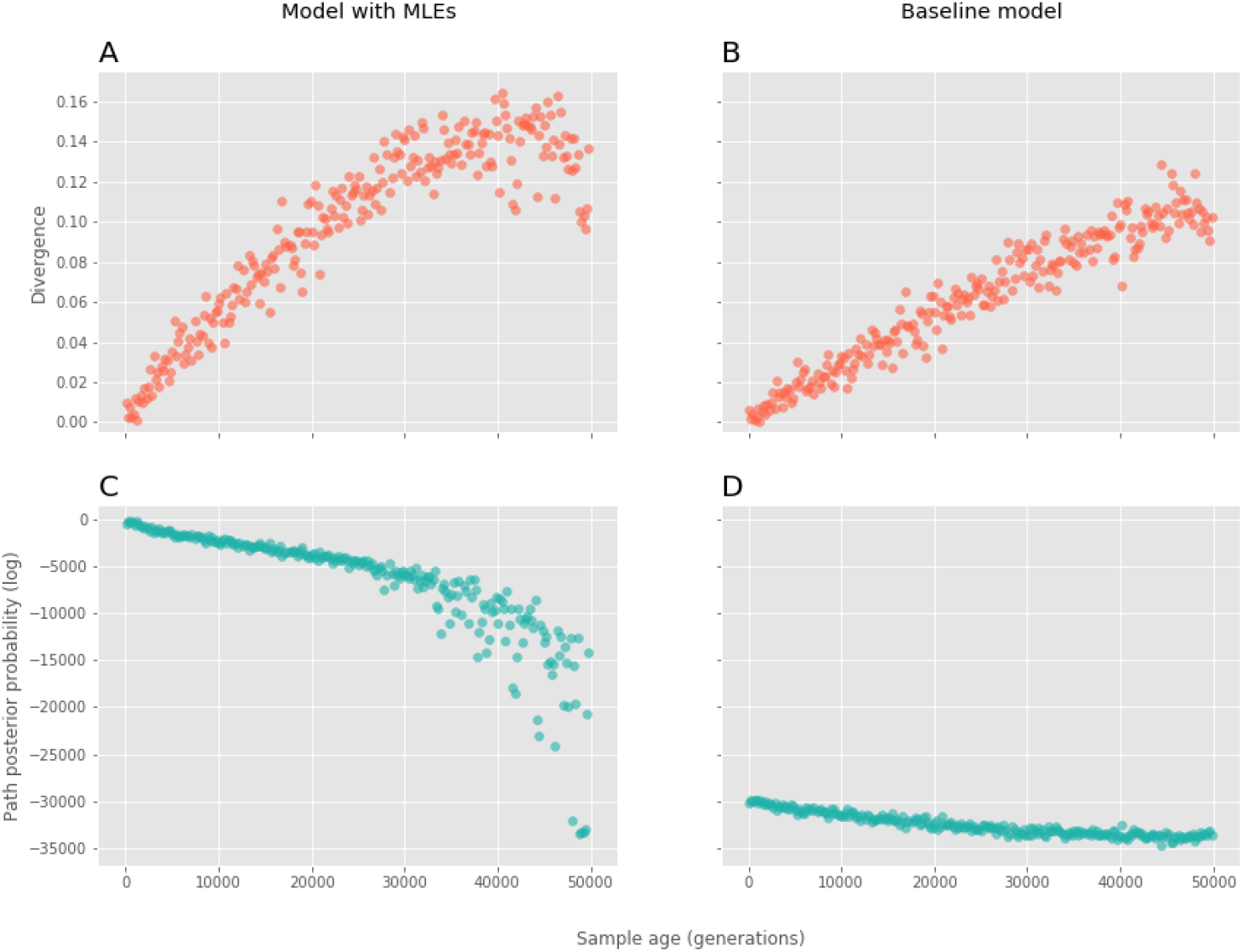
Comparison of the maximum likelihood (ML) model to a baseline model with uniform transition probabilities. **A** and **B** correspond to the divergence of the target haplotype from the optimal path against sample age for the ML model and the baseline model, respectively. **C** and **D** correspond to the log posterior probabilities of the optimal path against sample age for the ML model and the baseline model, respectively.

When looking at the log posterior probabilities of the optimal paths, we see that for the baseline model, they decrease linearly with sample age (i.e. the probabilities decrease exponentially with sample age), with values in the range of -30,000 to -35,000 (Figure 2d). For the ML model, there is a linear decrease in log probabilities (i.e. an exponential decrease in probability) as a function of sample age until about 30,000 generations ago, where the trend becomes exponential. It is worth noting that the values for some of the oldest samples reach the range of the baseline model (Figure 2c), indicating a drop in model performance.

### 3.3 Distribution of Divergences from the Reference Haplotypes

Upon comparison of the divergence from the optimal path with the mean divergence from the modern reference haplotypes, we see that the mean increases linearly with sample age, staying in the range of 15 to 30% divergence approximately. The divergence from the optimal path increases linearly until about 30,000 generations ago, with the rate of increase being higher than for the mean divergence. Starting close to 0% for the more recent samples and plateauing between 10 to 15% divergence, the optimal path values never reach the mean from the reference panel. In fact, a slight decrease can be observed after 40,000 generations (Figure 3).

**Figure 3:**
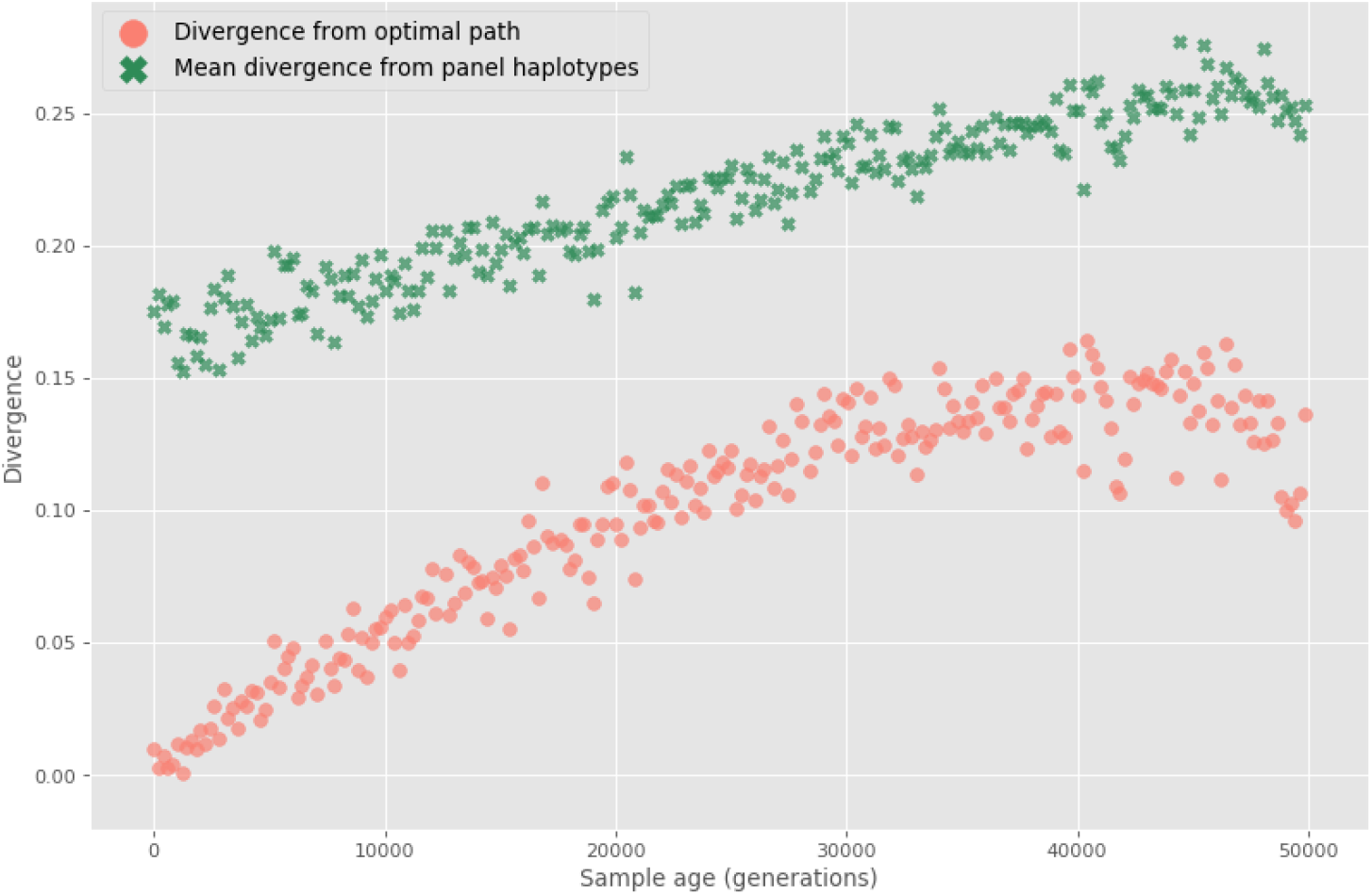
Comparison of the divergence from the optimal path and the mean divergence from reference to target haplotypes as a function of sample age. The red points correspond to the divergence from the optimal path to the target haplotype. The mean divergence from the reference panel is represented with green crosses.

Lastly, as can be seen in Figure 4, there is an exponential decrease in the variance of the divergence from reference to target haplotypes as a function of sample age. That is, the target haplotypes become equally divergent from all modern reference haplotypes at an exponential rate.

**Figure 4:**
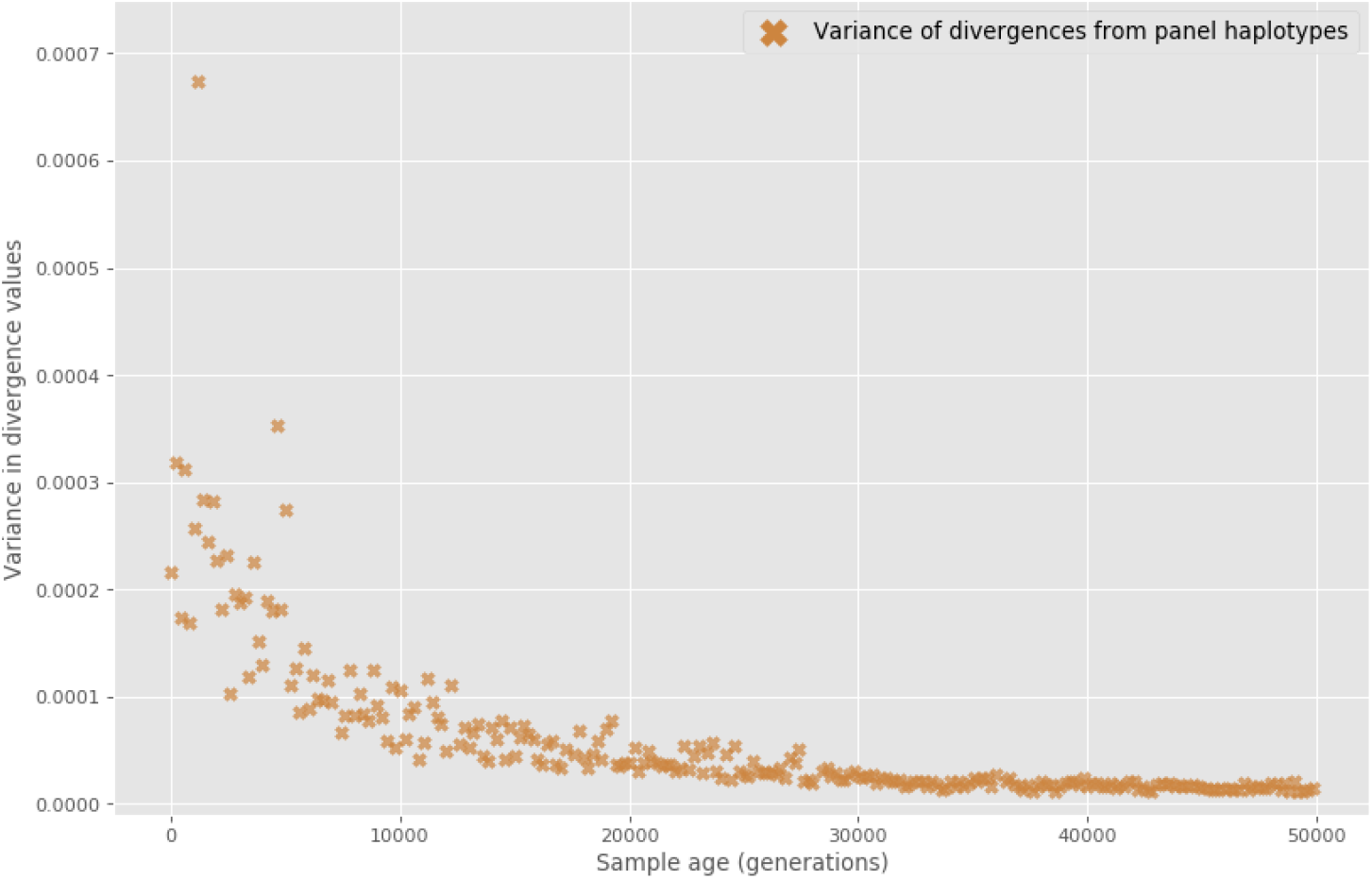
Variance in the divergence from reference to target haplotypes as a function of sample age.

## 4 Discussion

This study investigates the robustness of the haplotype-copying model when modeling ancient haplotypes as mosaics of modern haplotypes, motivated by new computational methods addressing challenges in aDNA analysis (Huang and Ringbauer, 2022; Ringbauer et al., 2021; Ausmees and Nettelblad, 2022). Biddanda *et al*. previously studied a haplotype-copying model with genetic distances assumed known (Biddanda et al., 2022). However, their approach had two main caveats: the maximum sample age was 15,000 years, limiting the analysis to a recent portion of human history, and the use of coalescent simulation to generate genetic variation data, which could result in lower overall genetic diversity and biased results (Gutenkunst et al., 2009; Yuan et al., 2012). To address these limitations, our study simulated a population starting 1.5M years ago using forward-time simulation.

In this 1.5 million-year-old population, at first, the jump rate increases monotonically with sample age, consistent with Biddanda *et al*.’s hypothesis (Biddanda et al., 2022). Then it becomes exponential past 30,000 generations. The model’s performance decreases as target haplotypes become equally divergent from reference haplotypes, resulting in high values of *λ* and low log posterior probabilities of optimal paths. The model’s effectiveness started to cease around 30,000 generations (900,000 years) ago.

Possible explanations for the model’s performance decline can be found in coalescent theory, which underlies coalescent simulation and dictates that except for very small samples, the coalescent time of a sample is close to that of the population (Rosenberg and Nordborg, 2002). Further, a significant portion of the coalescent time is spent on the last few lineages to coalesce (Gilean McVean, 2001). Thus, it is expected that most modern lineages will have coalesced going 30,000 generations back in time, making the time separation far too great for the model to identify genealogical relationships between the target and reference haplotypes.

Altogether, the findings of this study highlight the need for further investigation into the haplotype-copying model’s performance and limitations when dealing with ancient populations and aDNA analysis.

## 5 Conclusion

In conclusion, our study investigated the impact of ancient haplotype sampling on time separation with modern reference haplotypes, revealing the effects of coalescence and lineage variability. Although the model has limitations due to demographic assumptions and simplified genetic data, incorporating more realistic scenarios and data types could enhance its applicability. Future work should address these limitations to better understand shared ancestry between ancient and modern haplotypes.

## 6 Acknowledgements

The Novo Nordisk Data Science Investigator grant number NNF20OC0062491 provided the funding for this research project and the PhD scholarship of JDR. We would like to acknowledge the useful comments of Anders Albrechtsen. We would like to thank the Department of Health Tech at The Technical University of Denmark for additional funding and use of the HealthTech cluster.

## References

Kristiina Ausmees and Carl Nettelblad. Achieving improved accuracy for imputation of ancient DNA. bioRxiv, 2022. doi: 10.1101/2022.04.26.489533. URL https://www.biorxiv.org/content/early/2022/04/27/2022.04.26.489533x.

Franz Baumdicker, Gertjan Bisschop, Daniel Goldstein, Graham Gower, Aaron P Ragsdale, Georgia Tsambos, Sha Zhu, Bjarki Eldon, E Castedo Ellerman, Jared G Galloway, Ariella L Gladstein, Gregor Gorjanc, Bing Guo, Ben Jeffery, Warren W Kretzschumar, Konrad Lohse, Michael Matschiner, Dominic Nelson, Nathaniel S Pope, Consuelo D Quinto-Cortés, Murillo F Rodrigues, Kumar Saunack, Thibaut Sellinger, Kevin Thornton, Hugo van Kemenade, Anthony W Wohns, Yan Wong, Simon Gravel, Andrew D Kern, Jere Koskela, Peter L Ralph, and Jerome Kelleher. Efficient ancestry and mutation simulation with msprime 1.0. Genetics, 220(3):iyab229, 2022.

Arjun Biddanda, Matthias Steinrücken, and John Novembre. Properties of 2-locus genealogies and linkage disequilibrium in temporally structured samples. Genetics, 221, 3 2022. ISSN 1943-2631. doi: 10.1093/genetics/iyac038.

Gilean McVean. The coalescent. https://www.stats.ox.ac.uk/~mcvean/notes3.pdf, 2001. x[Online; accessed 2-August-2022].

Ryan N. Gutenkunst, Ryan D. Hernandez, Scott H. Williamson, and Carlos D. Bustamante. Inferring the Joint Demographic History of Multiple Populations from Multidimensional SNP Frequency Data. PLoS Genetics, 5:e1000695, 10 2009. ISSN 1553-7404. doi: 10.1371/journal.pgen.1000695.

Benjamin C. Haller, Jared Galloway, Jerome Kelleher, Philipp W. Messer, and Peter L. Ralph. Tree-sequence recording in SLiM opens new horizons for forward-time simulation of whole genomes. Molecular Ecology Resources, 19:552–566, 3 2019. ISSN 1755-098X. doi: 10.1111/1755-0998.12968.

Yilei Huang and Harald Ringbauer. hapCon: estimating contamination of ancient genomes by copying from reference haplotypes. Bioinformatics, 6 2022. ISSN 1367-4803. doi: 10.1093/bioinformatics/btac390.

Michael I. Jensen-Seaman, Terrence S. Furey, Bret A. Payseur, Yontao Lu, Krishna M. Roskin, Chin-Fu Chen, Michael A. Thomas, David Haussler, and Howard J. Jacob. Comparative Recombination Rates in the Rat, Mouse, and Human Genomes. Genome Research, 14:528–538, 4 2004. ISSN 1088-9051. doi: 10.1101/gr.1970304.

Jerome Kelleher, Yan Wong, Anthony W. Wohns, Chaimaa Fadil, Patrick K. Albers, and Gil McVean. Inferring whole-genome histories in large population datasets. Nature Genetics, 51:1330–1338, 9 2019. ISSN 1061-4036. doi: 10.1038/s41588-019-0483-y.

Daniel John Lawson, Garrett Hellenthal, Simon Myers, and Daniel Falush. Inference of Population Structure using Dense Haplotype Data. PLoS Genetics, 8:e1002453, 1 2012. ISSN 1553-7404. doi: 10.1371/journal.pgen.1002453.

Na Li and Matthew Stephens. Modeling Linkage Disequilibrium and Identifying Recombination Hotspots Using Single-Nucleotide Polymorphism Data. Genetics, 165:2213–2233, 12 2003. ISSN 1943-2631. doi: 10.1093/genetics/165.4.2213.

Martin Petr. slendr: A Simulation Framework for Spatiotemporal Population Genetics, 2022. URL https://github.com/bodkan/slendr. R package version 0.2.0.

Gabriel Renaud, Kristian Hanghøj, Thorfinn Sand Korneliussen, Eske Willerslev, and Ludovic Orlando. Joint Estimates of Heterozygosity and Runs of Homozygosity for Modern and Ancient Samples. Genetics, 212: 587–614, 7 2019. ISSN 1943-2631. doi: 10.1534/genetics.119.302057.

Harald Ringbauer, John Novembre, and Matthias Steinrücken. Parental relatedness through time revealed by runs of homozygosity in ancient DNA. Nature Communications, 12:5425, 12 2021. ISSN 2041-1723. doi: 10.1038/s41467-021-25289-w.

Noah A. Rosenberg and Magnus Nordborg. Genealogical trees, coalescent theory and the analysis of genetic polymorphisms. Nature Reviews Genetics, 3:380–390, 5 2002. ISSN 1471-0056. doi: 10.1038/nrg795.

Matthew Stephens, Nick J Smith, and Peter Donnelly. A new statistical method for haplotype reconstruction from population data. The American Journal of Human Genetics, 68(4):978–989, 2001.

Marc Tremblay and Héléne Vézina. New Estimates of Intergenerational Time Intervals for the Calculation of Age and Origins of Mutations. The American Journal of Human Genetics, 66:651–658, 2 2000. ISSN 00029297. doi: 10.1086/302770.

Xiguo Yuan, David J. Miller, Junying Zhang, David Herrington, and Yue Wang. An Overview of Population Genetic Data Simulation. Journal of Computational Biology, 19:42–54, 1 2012. ISSN 1066-5277. doi: 10.1089/cmb.2010.0188.

