## Supplemental Methods for "The Limits of Haplotype-Based Approaches: Exploring the Applicability of the Li and Stephens Haplotype-Copying Model to Ancient Samples"

June 21, 2023

### 1 Supplementary Methods

#### 1.1 Hidden Markov Models (HMMs)

Before diving into the specifics of the Li and Stephens model, some background information on HMMs needs to be provided. This section covers the basic theory of HMMs, including a formal definition and the statistical inference procedures used in this project.

##### 1.1.1 Formal Definition

HMMs are a class of statistical models that aim to capture unobservable information about the state of a system from a sequence of observable symbols. As such, an HMM consists of two stochastic processes: a series of underlying states hidden to the observer, and a sequence of observations whose probability distributions depend on the hidden states (Yoon, 2009).

According to Zucchini *et al.* (Zucchini et al., 2016), the hidden states in the first building block of an HMM form a Markov chain or process, formally defined as a sequence of discrete random variables $\{C_t : t \in \mathbb{N}\}$, that takes on values from a set of possible states $S = \{1, 2, \ldots, m\}$ and, for all $t \in \mathbb{N}$ [1], satisfies the Markov property:

$$\mathbb{P}(C_t|C_{t-1}, ..., C_1) = \mathbb{P}(C_t|C_{t-1}). \tag{1}$$

Meaning that the probability of being in a certain state at time $t$, only depends on the state the system was in at time $t - 1$[2].

The conditional probability of transitioning from state $i$ to state $j$ is known as the transition probability (Zucchini et al., 2016), denoted by

$$\gamma_{ij} = \mathbb{P}(C_t = j|C_{t-1} = i), \qquad i, j \in S. \tag{2}$$

The transition probabilities for the $m$ possible hidden states can be described by a transition matrix $\Gamma$, defined as the square matrix of dimensions $m \times m$ and row sums equal to 1 (Zucchini et al., 2016):

[1]This is true for a homogeneous Markov chain.
[2]This property holds for first-order Markov chains.

$$\Gamma = \begin{pmatrix} \gamma_{11} & \cdots & \gamma_{1m} \\ \vdots & \ddots & \vdots \\ \gamma_{m1} & \cdots & \gamma_{mm} \end{pmatrix}. \quad (3)$$

The unconditional probabilities of being in a certain state at time $t$ are denoted by the row vector $u(t) = (\mathbb{P}(C_t = 1), \ldots, \mathbb{P}(C_t = m)), \quad t \in \mathbb{N}$. Of particular interest is the initial distribution of the Markov process $u(1)$[3], oftentimes referred to in the literature as $\pi$ (Zucchini et al., 2016).

Zucchini *et al.* then go on to define the second building block of HMMs, i.e. the sequence of observed symbols $\{X_t : t \in \mathbb{N}\}$, in which the distribution of $X_t$ depends only on the current state $C_t$, hence

$$\mathbb{P}(X_t|X_1, X_2, \ldots, X_{t-1}, C_1, C_2, \ldots, C_t) = \mathbb{P}(X_t|C_t). \quad (4)$$

In this work, we are concerned with the case where the observations take on a value from a set of discrete symbols $O = \{O_1, O_2, \ldots, O_n\}$. As Zucchini *et al.* note, the probability of observing a certain symbol given that the system is in a certain state is called emission probability, denoted by

$$\sigma_i(x) = \mathbb{P}(X_t = x|C_t = i), \qquad x \in O, \quad i \in S. \quad (5)$$

Similarly to the transition probabilities, the emission probabilities can be stored in an emission matrix $\Sigma$, with row sums equal to 1, and dimensions $m \times n$, where $m$ is the number of possible hidden states and $n$ is the number of possible realizations of each observation:

$$\Sigma = \begin{pmatrix} \sigma_1(O_1) & \ldots & \sigma_1(O_n) \\ \vdots & \ddots & \vdots \\ \sigma_m(O_1) & \ldots & \sigma_m(O_n) \end{pmatrix}. \quad (6)$$

Figure 1 shows a schematic representation of the architecture of an HMM with two possible hidden states and an alphabet of three symbols which the observations can take values from.

[3]The initial distribution of the Markov chain $\pi$ will be used in the forward and the Viterbi algorithms, described in 1.1.3 and 1.1.4, respectively.

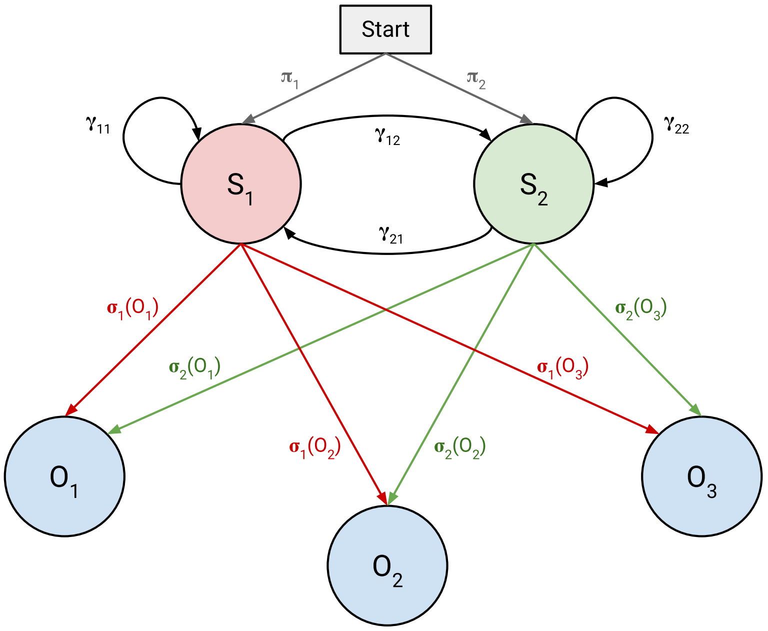

Figure 1: **Architecture of an HMM with two hidden states and three possible observation values.** In this example, the set of hidden states corresponds to $S = \{S_1, S_2\}$, and the observations can take on values from the alphabet $O = \{O_1, O_2, O_3\}$. The grey arrows denote the probabilities of starting the Markov process from either the red or the green hidden state, which are given by the initial distribution $\pi$. The transition probabilities are represented by the black arrows. As an example, $\gamma_{12}$ denotes the probability of transitioning from state $S_1$ to state $S_2$. Completing the architecture of the HMM, the red and green arrows denote the probabilities of emitting each possible observation symbol from the red ($S_1$) or the green state ($S_2$), respectively. For example, the probability of emitting symbol $O_1$ from state $S_2$, is given by $\sigma_2(O_1)$.

##### 1.1.2 Statistical Inference in HMMs

In a notorious tutorial on HMMs from 1989 (Rabiner, 1989), Rabiner established the notion that once the architecture of any HMM has been defined, there are three basic problems that characterize HMMs and that need to be solved in most settings where HMMs are used.

**Problem 1.** Given an HMM and a sequence of observations, what is the probability of observing that particular sequence?

**Problem 2.** Given an HMM and a sequence of observations, what is the most probable path of hidden states that generated that sequence?

**Problem 3.** Given a sequence of observations and a set of hidden states, what are the model parameters that best account for the data?

For clarity, the answers to these questions will be presented in the following subsections.

##### 1.1.3 The Forward Algorithm

This section answers the question posed by **Problem 1** (section 1.1.2), referred to as the evaluation problem by Rabiner, where a given model is evaluated in terms of the probability that a sequence of observations was produced by the model in question.

Formally, we wish to calculate the probability of a sequence of observed symbols $X = X_1, X_2, \ldots, X_T$ given the model $\lambda = (\Gamma, \Sigma, \pi)$, $\mathbb{P}(X|\lambda)$.

Rabiner begins by considering a specific sequence of hidden states $C = c_1, c_2, \ldots, c_T$.

The probability of observing the sequence of symbols $X$, assuming it was generated by the state sequence $C$ is given by

$$\mathbb{P}(X|C,\lambda) = \prod_{t=1}^{T} \mathbb{P}(X_t|c_t,\lambda) \tag{7}$$

or

$$\mathbb{P}(X|C,\lambda) = \sigma_{c_1}(X_1) \cdot \sigma_{c_2}(X_2) \ldots \sigma_{c_T}(X_T) \tag{8}$$

since an observation only depends on the current state of the system (Rabiner, 1989).

On the other hand, Rabiner defines the probability of the state sequence $C$ as

$$\mathbb{P}(C|\lambda) = \pi_{c_1}\gamma_{c_1c_2}\gamma_{c_2c_3}\cdots\gamma_{c_{T-1}c_T}. \tag{9}$$

The product of equations 8 and 9 gives us the probability of the observation sequence $X$ and the state sequence $C$ occurring simultaneously under the studied model (Rabiner, 1989):

$$\mathbb{P}(X,C|\lambda) = \mathbb{P}(X|C,\lambda)\mathbb{P}(C|\lambda). \tag{10}$$

By marginalizing over all possible sequences of hidden states, Rabiner obtains the probability of observing $X$ under the specified model $\lambda$:

$$\mathbb{P}(X|\lambda) = \sum_{C} \mathbb{P}(X|C,\lambda)\mathbb{P}(C|\lambda). \tag{11}$$

For an HMM with $m$ hidden states and an observation sequence of $T$ observations, there are $m^T$ possible state sequences, meaning that, although straightforward, computing probability 11 requires on the order of $m^T$ calculations. Even for relatively small values of $m$ and $T$ this task becomes computationally infeasible (Rabiner, 1989).

Luckily, a more efficient method that relies on dynamic programming exists, namely the forward algorithm. Rabiner starts by introducing the concept of forward probability $\alpha_t(i)$, defined as

$$\alpha_t(i) = \mathbb{P}(X_1, X_2, \ldots, X_t, c_t = S_i|\lambda). \tag{12}$$

This is the probability of the observation sequence $X$ until time $t$ and state $S_i$ at time $t$, given the model $\lambda$. As Rabiner describes, 12 can be solved recursively, by storing intermediate values for each partial observation sequence, in three steps:

1. Initialization:

$$\alpha_1(i) = \pi_i \sigma_i(X_1), \qquad 1 \leq i \leq m. \tag{13}$$

2. Recursion:

$$\alpha_{t+1}(j) = \big[\sum_{i=1}^{m} \alpha_t(i)\gamma_{ij}\big]\sigma_j(X_{t+1}), \qquad \begin{array}{l} 1 \leq t \leq T-1 \\ 1 \leq j \leq m. \end{array} \tag{14}$$

3. Termination:

$$\mathbb{P}(X|\lambda) = \sum_{i=1}^{m} \alpha_T(i) \tag{15}$$

In the first step, the forward probabilities are initialized as the joint probability of state $S_i$ and initial observation $X_1$. Next, in the recursion step (illustrated in Figure 2), the joint probability of emitting observation $X_{t+1}$ and being in state $S_j$ at time $t+1$ after seeing $X_1, X_2, \ldots, X_t$ is computed. The probability of the partial observation sequence up until time $t$ is stored in $\alpha_t(i)$ and the probability of arriving at state $S_j$ from state $S_i$ is given by $\gamma_{ij}$, thus by summing over the product of these two quantities over all the possible previous states $i$, the probability of the partial observation sequence up until time $t+1$ is obtained. Once we are in state $S_j$, the probability of observing $X_{t+1}$ is simply $\sigma_j(X_{t+1})$. Lastly, $\alpha_T(i)$ corresponds to the terminal forward variable, i.e. the probability of the entire observation sequence and ending up in state $S_i$. Therefore, the algorithm is terminated by summing $\alpha_T(i)$ over all possible hidden states, finally obtaining $\mathbb{P}(X|\lambda)$ (Rabiner, 1989).

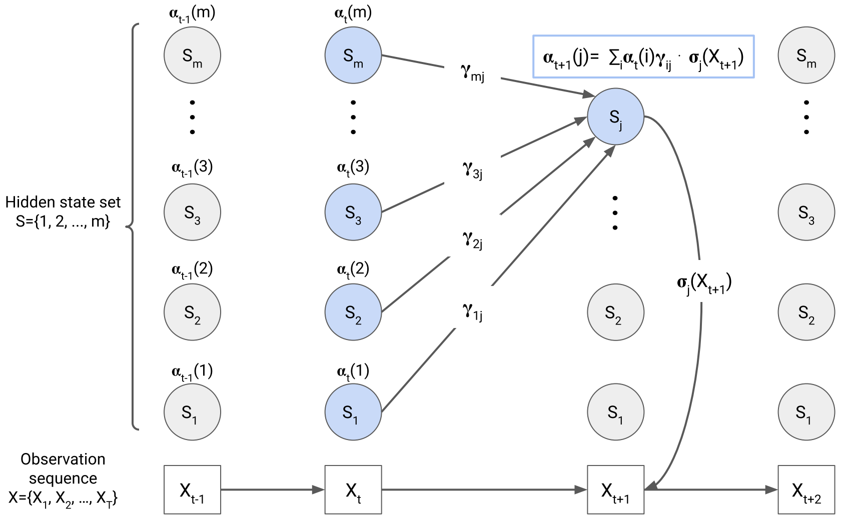

Figure 2: **Recursion step of the forward algorithm.** The forward probability $\alpha_{t+1}(j)$ is computed by summing all the previous forward probabilities ($\alpha_t$) weighted by their transition probabilities $\gamma$. This operation yields the probability of being in state $S_j$ at time $t+1$ after having seen the observation sequence $X_1, X_2, \ldots, X_t$. Lastly, the observation $X_{t+1}$ is accounted for by multiplying by the emission probability $\sigma_j(X_{t+1})$. Thus, we obtain the joint probability of the partial observation sequence $X_1, X_2, \ldots, X_{t+1}$ and being in state $S_j$ at time $t+1$. This diagram was adapted from (Daniel Jurafsky and James H. Martin, 2021).

With a time complexity of $O(m^2T)$, where $m$ is the number of states and $T$ the length of the observation sequence, the forward algorithm presents a far more efficient alternative to the brute-force computation of $\mathbb{P}(X|\lambda)$ (Rabiner, 1989).

##### 1.1.4 The Viterbi Algorithm

Since in most settings, HMMs are applied in order to infer information about the underlying properties of a system, it is oftentimes of interest to find the optimal sequence of hidden states, $C = c_1, c_1, \ldots, c_T$, that generated the observed data $X = X_1, X_2, \ldots, X_T$. That is, the state sequence that maximizes $\mathbb{P}(C|X, \lambda)$ (Rabiner, 1989). This is known as the decoding task (Brown and Golod, 2010), and it answers the question posed by **Problem 2** (section 1.1.2).

In order to find such a sequence, Rabiner begins by defining the variable

$$\delta_t(i) = \max_{c_1, c_2, \ldots, c_{t-1}} \mathbb{P}(c_1, c_2, \ldots, c_{t-1}, X_1, X_2, \ldots, X_t, c_t = i|\lambda) \tag{16}$$

which corresponds to the highest probability along a single path, at time $t$, that accounts for the first $t$ observations and ends in state $S_i$. Through recursion we get

$$\delta_{t+1}(j) = [\max_i \delta_t(i)\gamma_{ij}] \cdot \sigma_j(X_{t+1}). \tag{17}$$

In order to construct the state sequence, we have to keep track of the state that maximizes 17 at each timepoint $t$. For this we have an additional array $\psi_t(j)$ (Rabiner, 1989). As described by Rabiner, the algorithm, therefore, consists of the following steps:

1. Initialization:

$$\delta_1(i) = \pi_i \sigma_i(X_1), \qquad 1 \leq i \leq m \tag{18a}$$

$$\psi_1(i) = 0. \tag{18b}$$

2. Recursion:

$$\delta_t(j) = \max_{1\leq i\leq m} [\delta_{t-1}(i)\gamma_{ij}]\ \sigma_j(X_t), \qquad \begin{matrix} 2 \leq t \leq T \\ 1 \leq j \leq m \end{matrix} \tag{19a}$$

$$\psi_t(j) = \arg\max_{1\leq i\leq m} [\delta_{t-1}(i)\gamma_{ij}], \qquad \begin{matrix} 2 \leq t \leq T \\ 1 \leq j \leq m. \end{matrix} \tag{19b}$$

3. Termination:

$$\mathbb{P}^* = \max_{1\leq i\leq m} [\delta_T(i)] \tag{20a}$$

$$c_T^* = \arg\max_{1\leq i\leq m} [\delta_T(i)]. \tag{20b}$$

4. Backtracking:

$$c_t^* = \psi_{t+1}(c_{t+1}^*), \qquad t = T-1, T-2, \ldots, 1. \tag{21}$$

The initialization is similar to that of the forward algorithm, however during the recursion step, instead of summing over all possible states, we record the state which maximizes the probability of the observation sequence. When the recursion ends, 20a yields the posterior probability of the optimal hidden state path. Lastly, in the backtracking step, we make use of the backpointers stored in 19b to reconstruct such path (Rabiner, 1989).

##### 1.1.5 Parameter Estimation via Direct Maximization of the Likelihood

In sections 1.1.3 and 1.1.4, the model parameters were assumed known. However, oftentimes we will be in a situation where we have a new sequence of observations and we want to find the parameter set that maximizes its likelihood given a model architecture of interest (**Problem 3**, section 1.1.2). In the literature, this is referred to as the training task (Yoon, 2009), and although there is no optimal way of estimating the model parameters for any given finite observation sequence $X = X_1, X_2, \ldots, X_T$, we can choose the parameter set $\lambda = (\Gamma, \Sigma, \pi)$ that locally maximizes $\mathbb{P}(X|\lambda)$ through an iterative procedure (Rabiner, 1989).

Since we defined $\mathbb{P}(X|\lambda)$ as the sum of the terminal forward probabilities over all possible hidden states (section 1.1.3), the maximum likelihood estimates (MLEs) of the model parameters, i.e. the parameter values that maximize the likelihood of the data, correspond to those that maximize said sum (Zucchini et al., 2016).

#### 1.2 The Li and Stephens Haplotype-Copying Model

Biological data can, in many cases, be regarded as a sequence of discrete observations taking values from a finite alphabet of symbols (4 nucleotides in the case of nucleic acids and 20 amino acids in the case of proteins). Thus it is no surprise that after their popularity in the field of speech recognition in the 1980s (Levinson et al., 1983; Rabiner, 1989), HMMs began drawing the attention of molecular biologists a few years later (Churchill, 1989). By modeling protein or nucleic acid sequences as HMMs, information about underlying biological processes can be extracted. For this purpose, HMMs have been applied in a wide variety of scenarios, including alignment of DNA sequences (Yoon, 2009), detection of protein domains (Bateman and Chothia, 1996), gene finding (Krogh et al., 1994) or prediction of protein secondary structure (Asai et al., 1993).

Li and Stephens also exploited this property of biological sequence data when they applied an HMM to genetic variation data in order to elucidate patterns of Linkage Disequilibrium (LD)[4] across the human genome, with the particularity that the hidden states of the model consisted of other sequences (Li and Stephens, 2003).

In the LS model, the sequence of observed symbols corresponds to a haplotype, which is no other than the ordered genetic variants present in an individual's chromosome. Generally, when we speak of haplotypes, we refer to those sequences made up of genetic variants that are significantly present in a population, known as single-nucleotide polymorphisms (SNPs)

[4]Linkage disequilibrium (LD) is the non-independence, at a population level, of the alleles carried at different positions in the genome (Li and Stephens, 2003).

(Institute, 2022). Thus the information about the population can be inferred from such haplotypes. More precisely, Li and Stephens used their model to obtain estimates of the recombination rate in a human population (Li and Stephens, 2003).

##### 1.2.1 Recombination, Linkage Disequilibrium and Genetic Distance

Humans are eukaryotic diploid organisms, and their haploid gametes are produced in a specific type of cell division called meiosis. During meiosis, crossovers between the homologous pairs of maternal and parental chromosomes can occur, giving rise to the exchange of genetic material. This process is known as recombination, and it increases the genetic diversity in a population by ensuring that the offspring will inherit genetic information from each of its four grandparents (Clancy, 2008).

The further apart that two genes on the same chromosome are, the more likely they will undergo recombination. By the same token, this means that genetic variants that are physically close will tend to be inherited together. When two alleles are inherited together, we say they are linked, and when this phenomenon extends to the population level, we say the two variants are in linkage disequilibrium (LD) (Laird and Lange, 2011).

The concepts of linkage and LD are illustrated in Figure 3. A mutation occurs in the first generation. During meiosis, recombination takes place, and one of the chromosomes that the offspring inherits is a mosaic of the chromosome that contained the mutation and the one that did not. Some of these mosaic chromosomes will contain the mutation, and others will not. If the mutation is inherited, it will appear surrounded by the alleles that were present in the same parental chromosome, and thus we say they are linked. This is true only for those individuals in the population that are closely related, as it is unlikely that the same mutation occurred simultaneously in many families.

Many generations after, the mutation is present in a significant portion of the population. Subsequent recombination events have shortened the chromosomal segment that contained the mutation, but still, alleles that are physically close tend to appear together with the mutation. We say these alleles are in LD with the mutation. Thus, information about ancestral recombination events and recombination rates across the genome can be inferred by studying LD patterns in a population sample.

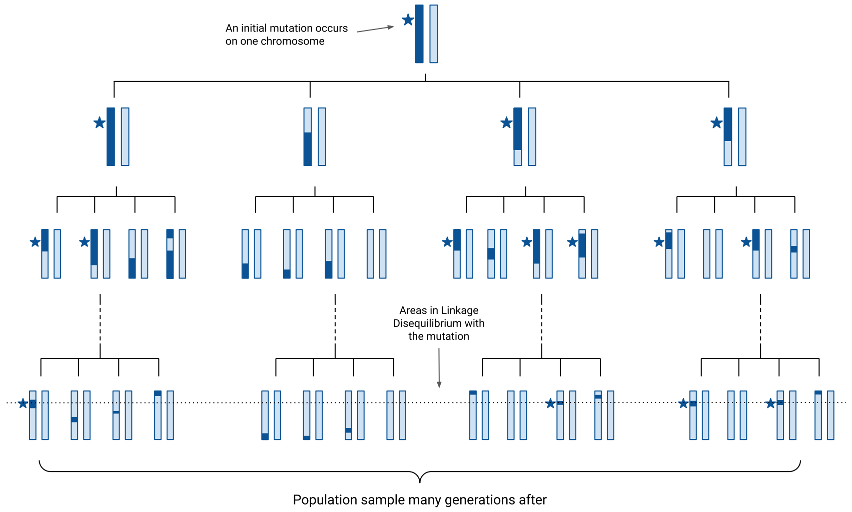

Figure 3: **Illustration of the concepts of linkage and Linkage Disequilibrium.** A mutation (denoted by a star) occurs on the dark chromosome in the first generation. The light chromosome carries a common allele. Due to recombination, the chromosomes transmitted to subsequent generations are mosaics of the two (dark and light) parental chromosomes. When the mutation is present, it appears surrounded by a chunk of the dark chromosomal background, shortened in subsequent generations due to crossover events. The first three rows represent closely related individuals and thus illustrate the concept of linkage. The last row shares a common ancestor many generations in the past and thus illustrates the concept of LD. This figure is adapted from (Laird and Lange, 2011).

Another essential concept in the LS model is that of genetic distance. While the physical distance between two SNPs measures how many base pairs apart they are located, the genetic distance measures the relative positions of two loci on a chromosome in terms of the expected number of crossovers between them (Laird and Lange, 2011). The unit of genetic distance is Morgans, where 1 Morgan corresponds to the distance between chromosome positions for which the average number of crossovers in a single generation is 1 (Shen and Gong, 2016).

##### 1.2.2 The Hidden Markov Model

In the model that Li and Stephens originally described, $k + 1$ haplotypes are sampled from a population. The first $k$ haplotypes will serve as the set of hidden states, from now on referred to as the reference panel, and the $k + 1_{th}$ haplotype is the observation sequence.

Each observation consists of a biallelic SNP, therefore it takes values from the alphabet $\{0, 1\}$, 0 and 1 meaning that the variant is absent or present at a given position in a haplotype, respectively.

The test or target haplotype, i.e. the observation sequence, can then be reconstructed as an imperfect mosaic of the other $k$ sampled haplotypes in the reference panel in a copying process which can be described by an HMM (Li and Stephens, 2003).

In this HMM, the initial probability distribution ($\pi$ in section 1.1.1) is given by

$$\mathbb{P}(X_1 = x) = 1/k, \quad x \in 1, 2, \ldots, k \tag{22}$$

where $X_1$ represents the reference haplotype from which the target haplotype copies off at the first SNP (Li and Stephens, 2003).

Li and Stephens captured the effects of recombination in the model's transition probabilities, defined as

$$\mathbb{P}(X_{j+1} = x'|X_j = x) = \begin{cases} e^{-\rho d_j/k} + \frac{1}{k}(1 - e^{-\rho d_j/k}) & \text{if} \quad x' = x; \\ \frac{1}{k}(1 - e^{-\rho d_j/k}) & \text{otherwise} \end{cases} \tag{23}$$

where $X_j$ is the haplotype that $h_{k+1}$ copies from at site $j$, $d_j$ is the physical distance in base pairs between SNPs at sites $j + 1$ and $j$, $k$ is the size of the reference panel, and $\rho$ is the population-scaled recombination parameter that Li and Stephens estimate. More precisely, $\rho = 4N_e c$, where $N_e$ corresponds to the effective diploid population size and $c$ to the recombination rate in average crossovers per base pair per generation.

From this definition, it follows that the probability of zero crossovers between sites $j$ and $j + 1$ is $e^{-\rho d_j/k}$ and the probability of at least one crossover is $1 - e^{-\rho d_j/k}$. Given recombination, there is a probability $1/k$ of recombining to any of the $k$ reference haplotypes, including the haplotype copied at the previous SNP (Cardin, 2006).

Further, recombination events happen according to a Poisson process with rate $\rho d_j/k$ (Cardin, 2006). As $\rho d_j$ decreases, the probability of copying from the same haplotype as in the previous SNP becomes much higher than the probability of switching to a different state, capturing the fact that the higher the genetic distance[5] between two adjacent SNPs, the more likely a recombination event is. Li and Stephens hypothesize that due to a decrease in the probability of seeing an entirely novel haplotype, the rate should decrease as the sample size $k$ increases. Thus, the division by $k$.

[5]As the physical distance $d_j$ is in base pairs and the recombination rate $c$ is in units of average crossovers per base pair per generation, $d_j \times c$ yields genetic distance in Morgans (average crossovers per generation).

In order to account for mutation, Li and Stephens allow errors in the copying process with probability $\hat{\theta}/(k + \hat{\theta})$. On the other hand, the probability of the copy being exact is $k/(k + \hat{\theta})$. Specifically, if $h_{i,j}$ denotes the allele (0 or 1) at site $j$ in haplotype $i$, then, given the copying process $X_1, \dots, X_S$, the alleles $h_{k+1,1}, h_{k+1,2}, \dots, h_{k+1,S}$ are independent, with

$$\mathbb{P}(h_{k+1,j} = a | X_j = x, h_1, \dots, h_k) = \begin{cases} \frac{k}{k+\hat{\theta}} + \frac{1}{2}\frac{\hat{\theta}}{k+\hat{\theta}} & , h_{x,j} = a \\ \frac{1}{2}\frac{\hat{\theta}}{k+\hat{\theta}} & , h_{x,j} \neq a \end{cases} \tag{24}$$

These are the emission probabilities of the model and they arise from the idea that if $h_1, \dots, h_{k+1}$ are uniformly sampled from a neutrally-evolving, randomly-mating, constant-sized population, the probability that $h_{k+1}$ is separated by at least one mutation from the other $k$ haplotypes at a given site is $\theta/(k + \theta)$, where $\theta = 4N\mu$, and $\mu$ is the mutation rate per base pair per generation and $N$ is the size of the population (Li and Stephens, 2003). Li and Stephens avoid estimating this population-scaled mutation rate and fix the value of $\hat{\theta}$ to

$$\hat{\theta} = (\sum_{m=1}^{n-1} \frac{1}{m})^{-1} \tag{25}$$

where $n$ is the number of sampled haplotypes.

Lastly, the factor 1/2 ensures that as $\hat{\theta}$ approaches infinity, both alleles are equally likely (Li and Stephens, 2003).

#### 1.3 Applications of the LS model

In this section, representative examples of the three possible relationships in time between the target and reference haplotypes are presented.

##### 1.3.1 Copying Modern from Modern

The LS model is a model for patterns of LD. As we have covered, LD arises from the shared history of DNA sequences. Therefore, many of its applications are aimed at gaining information about the ancestry of DNA sequences but differ in the approach.

One application that aims at identifying shared ancestry of haplotypes and characterizing substructure in a population is the tool `Chromopainter`.

`Chromopainter` aims to capture the most relevant genealogical information about a haplotype sample by building a co-ancestry matrix. First, considering a single test or target haplotype, the LS model, or 'painting algorithm' as the authors name it, finds its closest relative at each position of the genome amongst all the other haplotypes. That is, the algorithm paints the recipient haplotype from chunks of donor haplotypes (Figure 4c). Given that at some positions of the genome, there may be more than a single closest relative, the algorithm can produce alternative realizations of the copying process (Figure 4d). Additionally, the expectation that a given haplotype acted as a donor at each position over an infinite number of such paintings is computed (Figure 4e). This operation is repeated using each of the sample haplotypes as recipients and the rest as donors. The expected number of chunks that each test haplotype received from each donor is recorded in the co-ancestry matrix (Figure 4f) (Lawson et al., 2012).

The motivation for this approach is the assumption that the most recent genealogical events capture population substructure well enough. If again we consider a single haplotype, at each locus, there is a genealogical tree, whose complete information is captured by the time to the most recent common ancestor (MRCA) with each of the other sample haplotypes (Figure 4b). The structure of this tree varies along the genome due to ancestral recombination events (Figure 4a). The authors hypothesize that the most recent genealogical events, meaning finding the closest relative at each locus, is enough to characterize the current substructure of the population (Lawson et al., 2012).

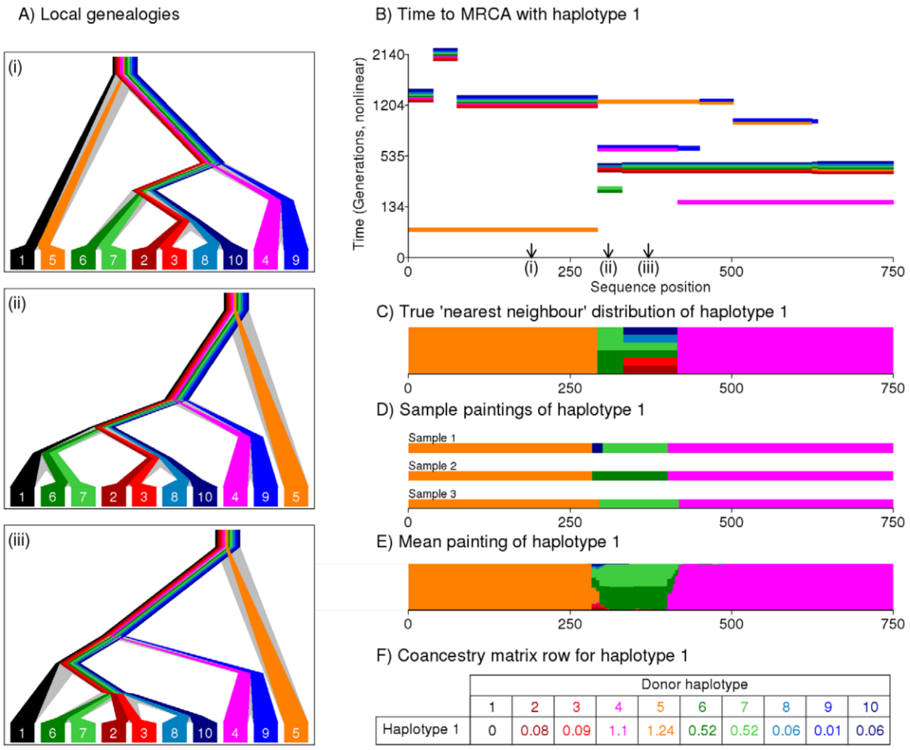

Figure 4: **Illustration of the painting process**. In this example, haplotype 1 acts as the recipient haplotype that is painted from chunks of all the other sample haplotypes. **A)** shows how the local genealogies vary at different loci, in **B)** the time to MRCA with haplotype 1 is shown at each position in the genome. In **C)** Haplotype 1 is painted from its closest relatives at each position. **D)** As in the middle of the sequence there are multiple closest relatives, the algorithm can produce different 'paintings'. The expectation of the painting process, estimating the closest relative distribution across the genomic positions is shown in **E)**. **F)** shows the row of the co-ancestry matrix for haplotype 1, derived from the expectation of the painting process. This figure is taken from the original Chromopainter publication (Lawson et al., 2012).

##### 1.3.2 Copying Modern from Ancient

Evolutionary biology focuses on inferring the genealogical history of DNA sequences in the hopes of shedding light on the events and forces that shape species and populations. As large genomic datasets are currently available, efficient inference methods are essential to exploit these resources fully. Tsinfer, a recently proposed method to overcome this efficiency issue, is based on the LS model. Starting from large datasets of modern genetic variation data, tsinfer deduces a set of haplotypes ancestral to those in the dataset. Then, using this set as a reference panel, the LS model is applied to a modern target haplotype, and the most probable path of hidden states (i.e. ancestral haplotypes) is inferred by employing the Viterbi algorithm thus reconstructing the genealogy of the modern query sequence (Kelleher et al., 2019).

Given that any extant haplotype must have predecessors moving back in time, the success of this method is guaranteed as long as the ancestral haplotypes are correctly inferred.

##### 1.3.3 Copying Ancient from Modern

Recent attempts at dealing with the issues covered in section 1.1.2 use a particular scenario of the LS model, where the target haplotype corresponds to an ancient individual, and the reference panel comprises modern haplotypes from present-day individuals.

One example is the tool `prophaser`. Although NGS technologies have increased the availability of ancient human genetic data, a large proportion of the available aDNA is sequenced at low coverage depths (1x or less). One way of maximizing the information obtained from sequencing such samples is the imputation of genotypes at missing sites or sites that can not be called confidently. Nevertheless, most genotype imputation tools are tailored to present-day genetic data, where studies are becoming increasingly large, and thus, computational efficiency is prioritized over accuracy. For this purpose, recent methods require phased genotypes as input. However, in aDNA data, it is not always possible to call genotypes confidently; thus, this approach may introduce errors (Ausmees and Nettelblad, 2022).

`prophaser` implements a diploid version of the LS model, where the observed data consists of genotype likelihoods instead of SNP data. First, using a reference panel of modern haplotypes, the posterior probabilities for the hidden states given the sequence of observed data at each position are calculated. Then, phased haplotypes are obtained by selecting the state with the highest probability and posterior genotype probabilities by integration over the hidden states. This way, both the phasing of the genotypes into haplotypes and the imputation of missing variants can be performed simultaneously (Ausmees and Nettelblad, 2022).

The results from the preprint show that `prophaser` outperforms two commonly used pipelines for phasing and imputation, particularly below 0.5x coverage depths. However, `prophaser` was tested on simulated data that consisted of modern genomes from the 1000 Genomes dataset (Auton et al., 2015), down-sampled in coverage, and on empirical data of 5 ancient individuals from around 3,000 to 9,000 years ago. Although the individuals used to simulate the low coverage data were then removed from the reference panel, one could argue that, even if down-sampled, the target haplotypes will not significantly differ from those in the reference panel, giving an overestimation of `prophaser`'s performance at low coverage depths. Additionally, the applicability of the tool to older genomes is not proven.

Another recently proposed tool based on the LS model where the target haplotype is ancient and the reference haplotypes are modern is `HapCon`. `HapCon` estimates contamination rates by applying the LS model to ancient male X chromosomes. By incorporating three error parameters that determine the emission probabilities of the model, the number of read counts that support either the reference or the alternative allele is modeled at each marker. The first parameter, $\epsilon_g$, corresponds to the error rate per base and aims to capture sequencing errors, characteristic aDNA damage and mismapping. Second, $\epsilon_r$ is the copying error from the original model, capturing mutation. Lastly, the parameter that `HapCon` estimates by maximum likelihood, $c$, corresponds to the contamination rate. Leveraging the fact that only mismatches due to contamination correlate with population allele frequencies, mismatches between the ancient target and the modern reference haplotypes are modeled either as errors or contamination (Huang and Ringbauer, 2022).

The contaminated sequences are simulated by mixing two BAM files from different individuals. The authors of `HapCon` report good performance on samples where the endogenous source is as old as 45,000 years. The endogenous source in this sample belongs to the Ust'-Ishim individual (Fu et al., 2014). Although it is one of the oldest modern human genomes ever sequenced, it is also significantly drifted with respect to present-day populations (Hawks, 2014). The authors state, 'When the genetic ancestry of contamination and endogenous sources are similar, the endogenous source can be closer to the allele frequencies of the specified contamination source than to the ones of the diverse reference panel [...]. And when the contaminant allele frequency is a better fit for the endogenous source than the reference panel, there is a bias toward the contamination source.' (Huang and Ringbauer, 2022). Conversely, a very drifted endogenous source means that the allele frequency of the contaminant source will be a much better fit for the reference panel, making the estimation of the contamination rate a more manageable task.

#### 1.4 Simulation of Genetic Variation Data at the Population Level

Population simulation plays an essential role in the field of evolutionary biology, allowing researchers to investigate how different demographic and genetic models affect patterns of genetic variation and evaluate the performance of various computational tools (Yuan et al., 2012). In the context of this project, it will allow us to test the performance of the LS model when reconstructing an ancient haplotype as an imperfect mosaic of modern reference haplotypes.

The two most used approaches to the simulation of population data are backward-time or coalescent simulation and forward or forward-time simulation, each with its strengths and weaknesses.

In a coalescent simulation, a sample of individuals representing the present generation is drawn from an idealized population with the desired characteristics. Then, the genealogical history of the sample is traced back in time until all lineages have coalesced[6], thus reaching the MRCA to all individuals (Yuan et al., 2012). Hence, any ancient haplotype sampled from this population will have direct descendants in the present generation.

On the other hand, forward simulation starts from an initial population in the past and then traces its evolution under the specified demographic scenario over the desired number of generations (Yuan et al., 2012). Typically, coalescent simulation is then performed, using the initial set of individuals as the starting point, in order to provide a prior genealogical history for any genomic segments that have not yet coalesced. This procedure is known as recapitation (Haller et al., 2019).

The general architecture of both coalescent and forward simulation is illustrated in Figure 5.

[6]A coalescence event is a point in a genealogy where two lineages meet into a common ancestor.

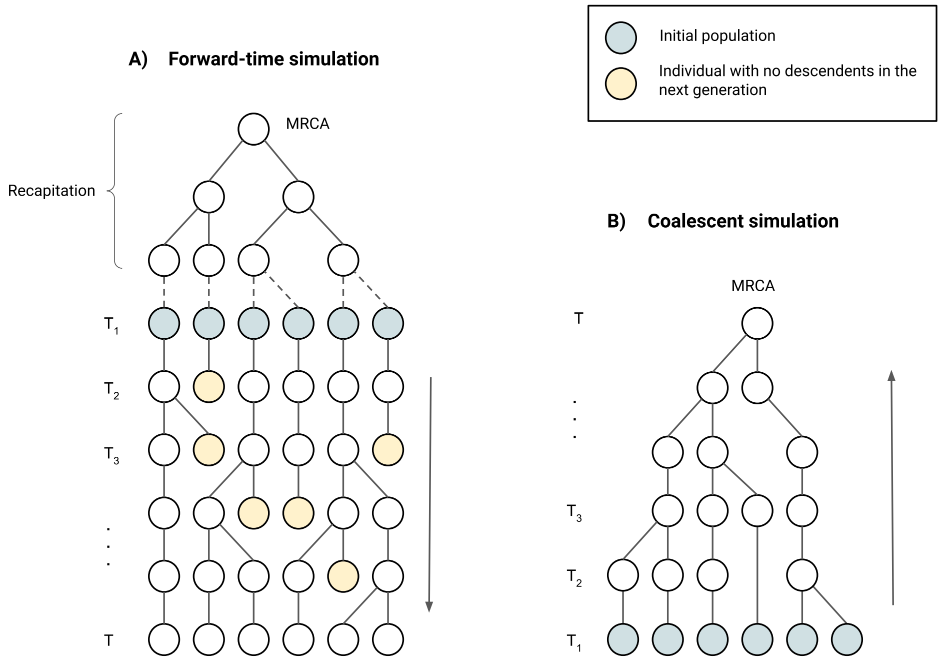

Figure 5: **Schematic representation of two populations simulated through the forward-time and the coalescent approach. A** shows the mechanistic of forward-time simulation, where starting from a set of individuals at generation $T_1$, which in this case is in the past, the entire history of the population is simulated for $T$ generations. Forward-time simulation allows the sampling of individuals that did not have any descendants, represented in yellow. Through recapitation, the MRCA to the initial population is reached. In **B** the genealogical history of a sample of individuals at time $T_1$, the present generation in this case, is simulated through the coalescent approach. The MRCA to all individuals in the present generation is reached at time $T$. In both **A** and **B**, the initial population is shown in blue.

Since forward simulation allows us to track the entire ancestral information of the population, it is computationally heavier than coalescent simulation, where we only have access to the genealogical history of the individuals in the present generation (Yuan et al., 2012). However, as an ancient sample from which DNA has been extracted might not have descendants among modern individuals, forward simulation provides a more realistic scenario to test the LS model's applicability.

Furthermore, as shown in Figure 5, a population simulated through coalescent simulation is much shallower than one that has been simulated through forward simulation and then recapitated. Hence, even if we were to perform coalescent simulation and then sample an individual who does not have direct descendants included in the modern reference panel, the MRCA to all modern individuals would be much closer in time than that of a forward simulation and the divergence between modern and ancient haplotypes would be lower on average. Thus, to avoid overestimating the LS model's performance, a forward simulation of genetic variation data at the population level was chosen.

##### 1.4.1 Forward-Time Simulations

**Demographic Model** In order to be able to isolate the effects of mutation and recombination on the performance of the haplotype-copying model as sample age increases, the simulations were performed under a simple demographic model consisting of a single continuous population with constant population size, neutral mutation and random mating. Two different populations were simulated. The first is intended to emulate the African population of anatomically and genetically modern humans that is widely accepted to have emerged around 200,000 years ago (Gutenkunst et al., 2009), and so this was the starting time for the simulation.

In light of the results obtained for the African population, it was deemed necessary to go further back in time to dissect the behavior of the haplotype-copying model. Thus, another population starting 1.5 million years ago was simulated. The rest of the parameters were kept the same as for the African population.

As previously established in the literature, generation time for modern humans is of 30 years (Tremblay and Vézina, 2000), the recombination rate corresponds to $10^{-8}$ crossovers per base pair per generation (Jensen-Seaman et al., 2004), and the mutation rate is $2 \times 10^{-8}$ per base pair per generation (Renaud et al., 2019).

In order to choose a value for the population size parameter, Watterson's estimator of genetic diversity, $\theta = 4N_e\mu$, was used, where $\mu$ corresponds to the mutation rate and $N_e$ corresponds to effective population size. Given that the value of $\theta$ for the African population has been empirically determined to be around $1.15 \times 10^{-3}$ (Renaud et al., 2019), an effective population size of 14,375 is obtained[7].

Lastly, the simulated sequence was 1Mb long.

[7] The effective population size $N_e$ is the size that an abstract, idealized population would need to be to match a given value of a particular metric. That value is typically estimated from an actual population (Haller and Messer, 2016). In our case, it corresponds to the African population's empirical estimate of $\theta$. Since our simulated populations are ideal populations with constant size, random mating and no selection, the chosen size corresponds to $N_e$.

**Simulation Engines** Genomic data at the population level was simulated through the `slendr` package (Petr, 2022) in `R`, choosing the forward-time simulator `SLiM` (Haller et al., 2019) as a back-end simulation engine.

The output of `SLiM` consists of tree-sequence files, a format that allows the compact storing of the population's genealogical information and genotypes (Haller et al., 2019).

Recapitulation of the tree-sequences was performed in `slendr`, using `msprime` (Baumdicker et al., 2022) as the coalescent simulation engine.

##### 1.4.2 Haplotype-Copying Model

The haplotype-copying model used in this work is a version of the original LS model, implemented and described by Biddanda et al. (Biddanda et al., 2022). The main difference with the original model is that the genetic distance between adjacent SNPs is assumed known. Li and Stephens estimate the population-scaled recombination parameter $\rho = 4N_ec$, where $N_e$ corresponds to the effective diploid population size, and $c$ corresponds to the recombination rate (Li and Stephens, 2003). In the work by Biddanda et al., the recombination parameter is assumed known and set to $10^{-8}$ crossovers per base pair per generation. As the physical distance between adjacent SNPs is also known, the genetic distance in Morgans can then be calculated as $g_l = c \times d_l$, where $g_l$ and $d_l$ are the genetic and physical distances, respectively, between the SNPs at sites $l$ and $l-1$.

The transition probabilities between the haplotypes in the modern reference panel, denoted $h_1, ..., h_k$, are then defined as follows:

$$P(X_l = x' | X_{l-1} = x) = \begin{cases} e^{-\lambda g_l} + \frac{1}{k}(1 - e^{-\lambda g_l}) & , x' = x \\ \frac{1}{k}(1 - e^{-\lambda g_l}) & , \text{else} \end{cases} \tag{26}$$

where $X_l$ denotes the haplotype from the reference panel that is being copied at site $l$, $k$ corresponds to the reference panel size and $g_l$ corresponds to the aforementioned genetic distance (Biddanda et al., 2022). Thus, there is an equal probability of jumping to any other haplotype in the reference panel and this probability increases exponentially with the genetic distance between the previous and the current SNP. Similarly, the probability of staying in the current state decreases exponentially as a function of the genetic distance. The $\lambda$ parameter, which Biddanda *et al.* estimate, determines the rate at which the probability of jumping to any other state increases with the genetic distance, and thus it can be interpreted as a jump rate parameter in units of the average number of jumps per Morgan (Biddanda et al., 2022). The behavior of the transition probabilities as a function of genetic distance and for different values of $\lambda$ is illustrated in Figure 1 in the main document.

In order to account for mutation, errors in the copying process are allowed as defined in the model's emission probabilities:

$$P(h_l = a | X_l = x) = \begin{cases} \epsilon & , h_{xl} \neq a \\ (1-\epsilon) & , h_{xl} = a \end{cases} \tag{27}$$

where $h_l$ and $h_{xl}$ refer to the allelic states (0 or 1) of the test and copied haplotypes at site $l$, respectively, and $\epsilon$ is the copying error parameter (Biddanda et al., 2022).

Lastly, Biddanda *et al.* use a uniform initial probability distribution, where the probability of starting the copying process from any given haplotype is $1/k$.

##### 1.4.3 Datasets

A reference panel of size $k = 100$ modern haplotypes was randomly selected for each of the populations.

Target haplotypes were sampled uniformly every 100 generations for the African population and every 200 generations for the 1,5 million-year-old population.

##### 1.4.4 Maximum Likelihood Estimation of Model parameters

MLEs of the jump rate and copying error parameters denoted $\hat{\lambda}$ and $\hat{\epsilon}$, respectively, were obtained via direct maximization of the likelihood (see section 1.1.5). The implementation belongs to Biddanda et al. (Biddanda et al., 2022). First, $\epsilon$ is fixed to $10^{-2}$ and profile MLEs of $\lambda$ are obtained by numerically minimizing the negative log likelihood using `scipy.optimize.minimize_scalar` (Virtanen et al., 2020). Then, $\hat{\lambda}$ and $\hat{\epsilon}$ are jointly estimated by two-dimensional numerical minimization of the negative log-likelihood function, using `scipy.optimize.minimize` (Virtanen et al., 2020) and the profile MLE of $\lambda$ and $\epsilon = 10^{-2}$ as initial values for the parameter space search. Finally, following Biddanda et al.'s implementation, standard errors for the MLEs were obtained via a finite-difference approximation to the second derivative of the joint log-likelihood surface.

##### 1.4.5 Viterbi Decoding

Once MLEs of the jump rate and copying error parameters were obtained, the most probable path of hidden states and its log posterior probability were computed for each test haplotype via the Viterbi algorithm (section 1.1.4).

Viterbi decoding was also applied to the data using a baseline model, where all the transition probabilities are equal to $1/k$. This means that the probability of jumping to any given state and of staying in the same state are equal. The value of the copying error parameter in this model was still $\hat{\epsilon}$.

The reason for the posterior probabilities of the optimal paths being in the log scale is that the intermediate path probabilities of the Viterbi algorithm are computed in log space in order to prevent numerical underflow due to the multiplication of small values. Hence, the log probabilities are summed according to the logarithm property

$$\log xy = \log x + \log y \quad (x, y > 0) \tag{28}$$

The Viterbi algorithm was implemented from scratch.

##### 1.4.6 Model Evaluation Metrics

Three different metrics were employed to evaluate the model's performance as the sample age increases.

The first metric is the MLE of the jump rate parameter, denoted $\hat{\lambda}$. Biddanda *et al.* proposed $\hat{\lambda}$ as a proxy for the model's accuracy if it were to be used in the context of missing data imputation (Biddanda et al., 2022). $\hat{\lambda}$ captures the average number of jumps per Morgan that occur in the copying process, and thus it is expected to increase with sample age, as the longer the time separation between the modern and ancient haplotypes is, the more recombination events can take place.

In addition to $\hat{\lambda}$, two other metrics were used: the divergence between the target haplotype and the optimal path of hidden states and the log posterior probability of this path. The optimal path corresponds to the mosaic of modern reference haplotypes with the highest posterior probability given the model and the observed data. First, it is obtained via the Viterbi algorithm, and then divergence is calculated as follows:

$$\text{divergence} = \frac{\text{edit distance}}{\text{total number of SNPs}} \tag{29}$$

where the edit distance corresponds to the number of mismatches between the two haplotypes. Normalizing by the total number of SNPs guarantees this metric to be comparable between haplotypes of different lengths.

The divergence from the optimal path and the optimal path's log posterior probability obtained under the maximum likelihood (ML) model, i.e. the model that maximizes the likelihood of the data, were then compared to those obtained under the baseline model described in the previous section (1.4.5).

Finally, we looked for changes in the distribution of divergences from the modern reference haplotypes to the target haplotypes as a function of their age, hoping to find an explanation for the model's behavior as the target haplotypes' age increases.

##### 1.4.7 Code

The code and data used in this project can be found in the GitHub repository `https://github.com/isadpc/HapCopying`.

#### References

Kiyoshi Asai, Satoru Hayamizu, and Ken'ichi Handa. Prediction of protein secondary structure by the hidden Markov model. *Bioinformatics*, 9:141–146, 1993. ISSN 1367-4803. doi: 10.1093/bioinformatics/9.2.141.

Kristiina Ausmees and Carl Nettelblad. Achieving improved accuracy for imputation of ancient DNA. *bioRxiv*, 2022. doi: 10.1101/2022.04.26.489533. URL `https://www.biorxiv.org/content/early/2022/04/27/2022.04.26.489533`.

Adam Auton et al. A global reference for human genetic variation. *Nature*, 526:68–74, 10 2015. ISSN 0028-0836. doi: 10.1038/nature15393.

Alex Bateman and Cyrus Chothia. Fibronectin type III domains in yeast detected by a hidden Markov model. *Current Biology*, 6:1544–1547, 12 1996. ISSN 09609822. doi: 10.1016/S0960-9822(02)70765-3.

Franz Baumdicker, Gertjan Bisschop, Daniel Goldstein, Graham Gower, Aaron P Ragsdale, Georgia Tsambos, Sha Zhu, Bjarki Eldon, E Castedo Ellerman, Jared G Galloway, Ariella L Gladstein, Gregor Gorjanc, Bing Guo, Ben Jeffery, Warren W Kretzschumar, Konrad Lohse, Michael Matschiner, Dominic Nelson, Nathaniel S Pope, Consuelo D Quinto-Cortés, Murillo F Rodrigues, Kumar Saunack, Thibaut Sellinger, Kevin Thornton, Hugo van Kemenade, Anthony W Wohns, Yan Wong, Simon Gravel, Andrew D Kern, Jere Koskela, Peter L Ralph, and Jerome Kelleher. Efficient ancestry and mutation simulation with msprime 1.0. *Genetics*, 220(3):iyab229, 2022.

Arjun Biddanda, Matthias Steinrücken, and John Novembre. Properties of 2-locus genealogies and linkage disequilibrium in temporally structured samples. *Genetics*, 221, 3 2022. ISSN 1943-2631. doi: 10.1093/genetics/iyac038.

Daniel G Brown and Daniil Golod. Decoding HMMs using the k best paths: algorithms and applications. *BMC Bioinformatics*, 11:S28, 1 2010. ISSN 1471-2105. doi: 10.1186/1471-2105-11-S1-S28.

Niall Cardin. *Approximating the Coalescent with Recombination.* PhD thesis, Corpus Christi College, University of Oxford, 2006.

Gary A. Churchill. Stochastic models for heterogeneous DNA sequences. *Bulletin of Mathematical Biology*, 51:79–94, 1 1989. ISSN 0092-8240. doi: 10.1007/BF02458837.

Suzanne Clancy. Genetic Recombination. *Nature Education*, 1(1): 40, 2008. URL `https://www.nature.com/scitable/topicpage/genetic-recombination-514/`. [Online; accessed 4-August-2022].

Daniel Jurafsky and James H. Martin. Speech and Language Processing. Chapter A: Hidden Markov Models. `https://web.stanford.edu/~jurafsky/slp3/A.pdf`, 2021. [Online; accessed 4-August-2022].

Qiaomei Fu et al. Genome sequence of a 45,000-year-old modern human from western Siberia. *Nature*, 514:445–449, 10 2014. ISSN 0028-0836. doi: 10.1038/nature13810.

Ryan N. Gutenkunst, Ryan D. Hernandez, Scott H. Williamson, and Carlos D. Bustamante. Inferring the Joint Demographic History of Multiple Populations from Multidimensional SNP Frequency Data. *PLoS Genetics*, 5:e1000695, 10 2009. ISSN 1553-7404. doi: 10.1371/journal.pgen.1000695.

B.C. Haller and P.W. Messer. SLiM: An Evolutionary Simulation Framework. `http://benhaller.com/slim/SLiM_Manual.pdf`, 2016. [Online; accessed 6-August-2022].

Benjamin C. Haller, Jared Galloway, Jerome Kelleher, Philipp W. Messer, and Peter L. Ralph. Tree-sequence recording in SLiM opens new horizons for forward-time simulation of whole genomes. *Molecular Ecology Resources*, 19:552–566, 3 2019. ISSN 1755-098X. doi: 10.1111/1755-0998.12968.

John Hawks. The genome from Ust´-Ishim. `https://johnhawks.net/weblog/reviews/ancient-genomes/ust-ishim-fu-2014.html`, 2014. [Online; accessed 5-August-2022].

Yilei Huang and Harald Ringbauer. hapCon: estimating contamination of ancient genomes by copying from reference haplotypes. *Bioinformatics*, 6 2022. ISSN 1367-4803. doi: 10.1093/bioinformatics/btac390.

National Human Genome Research Institute. Haplotype. `https://www.genome.gov/genetics-glossary/haplotype`, 2022. [Online; accessed 4-August-2022].

Michael I. Jensen-Seaman, Terrence S. Furey, Bret A. Payseur, Yontao Lu, Krishna M. Roskin, Chin-Fu Chen, Michael A. Thomas, David Haussler, and Howard J. Jacob. Comparative Recombination Rates in the Rat, Mouse, and Human Genomes. *Genome Research*, 14:528–538, 4 2004. ISSN 1088-9051. doi: 10.1101/gr.1970304.

Jerome Kelleher, Yan Wong, Anthony W. Wohns, Chaimaa Fadil, Patrick K. Albers, and Gil McVean. Inferring whole-genome histories in large population datasets. *Nature Genetics*, 51:1330–1338, 9 2019. ISSN 1061-4036. doi: 10.1038/s41588-019-0483-y.

Anders Krogh, I. Saira Mian, and David Haussler. A hidden Markov model that finds genes in E.coli DNA. *Nucleic Acids Research*, 22:4768–4778, 11 1994. ISSN 0305-1048. doi: 10.1093/nar/22.22.4768.

Nan M. Laird and Christoph Lange. *The Fundamentals of Modern Statistical Genetics*. Springer New York, 1 edition, 2011. ISBN 978-1-4419-7337-5. doi: 10.1007/978-1-4419-7338-2.

Daniel John Lawson, Garrett Hellenthal, Simon Myers, and Daniel Falush. Inference of Population Structure using Dense Haplotype Data. *PLoS Genetics*, 8:e1002453, 1 2012. ISSN 1553-7404. doi: 10.1371/journal.pgen.1002453.

S. E. Levinson, L. R. Rabiner, and M. M. Sondhi. An Introduction to the Application of the Theory of Probabilistic Functions of a Markov Process to Automatic Speech Recognition. *Bell System Technical Journal*, 62:1035–1074, 4 1983. ISSN 00058580. doi: 10.1002/j.1538-7305.1983.tb03114.x.

Na Li and Matthew Stephens. Modeling Linkage Disequilibrium and Identifying Recombination Hotspots Using Single-Nucleotide Polymorphism Data. *Genetics*, 165:2213–2233, 12 2003. ISSN 1943-2631. doi: 10.1093/genetics/165.4.2213.

Martin Petr. *slendr: A Simulation Framework for Spatiotemporal Population Genetics*, 2022. URL `https://github.com/bodkan/slendr`. R package version 0.2.0.

L.R. Rabiner. A tutorial on hidden Markov models and selected applications in speech recognition. *Proceedings of the IEEE*, 77:257–286, 1989. ISSN 00189219. doi: 10.1109/5.18626.

Gabriel Renaud, Kristian Hanghøj, Thorfinn Sand Korneliussen, Eske Willerslev, and Ludovic Orlando. Joint Estimates of Heterozygosity and Runs of Homozygosity for Modern and Ancient Samples. *Genetics*, 212: 587–614, 7 2019. ISSN 1943-2631. doi: 10.1534/genetics.119.302057.

Yiping Shen and Xiaohong Gong. Chapter 1 - Experimental Tools for the Identification of Specific Genes in Autism Spectrum Disorders and Intellectual Disability. In Carlo Sala and Chiara Verpelli, editors, *Neuronal and Synaptic Dysfunction in Autism Spectrum Disorder and Intellectual Disability*, pages 3–12. Academic Press, San Diego, 2016. ISBN 978-0-12-800109-7. doi: https://doi.org/10.1016/B978-0-12-800109-7.00001-7. URL `https://www.sciencedirect.com/science/article/pii/B9780128001097000017`.

Marc Tremblay and Hélène Vézina. New Estimates of Intergenerational Time Intervals for the Calculation of Age and Origins of Mutations. *The American Journal of Human Genetics*, 66:651–658, 2 2000. ISSN 00029297. doi: 10.1086/302770.

Pauli Virtanen, Ralf Gommers, Travis E. Oliphant, Matt Haberland, Tyler Reddy, David Cournapeau, Evgeni Burovski, Pearu Peterson, Warren Weckesser, Jonathan Bright, Stéfan J. van der Walt, Matthew Brett, Joshua Wilson, K. Jarrod Millman, Nikolay Mayorov, Andrew R. J. Nelson, Eric Jones, Robert Kern, Eric Larson, C J Carey, İlhan Polat, Yu Feng, Eric W. Moore, Jake VanderPlas, Denis Laxalde, Josef Perktold, Robert Cimrman, Ian Henriksen, E. A. Quintero, Charles R. Harris, Anne M. Archibald, Antônio H. Ribeiro, Fabian Pedregosa, Paul van Mulbregt, and SciPy 1.0 Contributors. SciPy 1.0: Fundamental Algorithms for Scientific Computing in Python. *Nature Methods*, 17:261–272, 2020. doi: 10.1038/s41592-019-0686-2.

Byung-Jun Yoon. Hidden markov models and their applications in biological sequence analysis. *Current Genomics*, 10:402–415, 9 2009. ISSN 13892029. doi: 10.2174/138920209789177575.

Xiguo Yuan, David J. Miller, Junying Zhang, David Herrington, and Yue Wang. An Overview of Population Genetic Data Simulation. *Journal of Computational Biology*, 19:42–54, 1 2012. ISSN 1066-5277. doi: 10.1089/cmb.2010.0188.

Walter Zucchini, Iain L. MacDonald, and Roland Langrock. *Hidden Markov Models for Time Series: An Introduction Using R*. Chapman and Hall/CRC, 2nd edition, 7 2016. ISBN 9781315372488. doi: 10.1201/b20790.
